# Neural markers of category-based selective working memory in aging

**DOI:** 10.1101/435388

**Authors:** Robert M. Mok, M. Clare O’Donoguhue, Nicholas E. Myers, Erin H.S. Drazich, Anna Christina Nobre

## Abstract

Working memory (WM) is essential for normal cognitive function, but shows marked decline in aging. Studies have shown that the ability to attend selectively to relevant information amongst competing distractors is related to WM capacity. The extent to which WM deficits in aging are related to impairments in selective attention is unclear. To investigate the neural mechanisms supporting selective attention in WM in aging, we tested a large group of older adults using functional magnetic resonance imaging whilst they performed a category-based (faces/houses) selective-WM task. Older adults were able to use attention to encode targets and suppress distractors to reach high levels of task performance. A subsequent, surprise recognition-memory task showed strong consequences of selective attention. Attended items in the relevant category were recognised significantly better than items in the ignored category. Neural measures also showed reliable markers of selective attention during WM. Purported control regions including the dorsolateral and inferior prefrontal and anterior cingulate cortex were reliably recruited for attention to both categories. Activation levels in category-sensitive visual cortex showed reliable modulation according to attentional demands, and positively correlated with subsequent memory measures of attention and WM span. Psychophysiological interaction analyses showed that activity in category-sensitive areas were coupled with non-sensory cortex known to be involved in cognitive control and memory processing, including regions in the PFC and hippocampus. In summary, we found that brain mechanisms of attention for selective WM are relatively preserved in aging, and individual differences in these abilities corresponded to the degree of attention-related modulation in the brain.

**Highlights:**

- Preserved attention for category-based selective working memory (WM) in aging
- Category-selective attention leads to improved recognition memory in aging
- Reliable top-down attention modulation in category-sensitive cortex in older adults
- Modulation of category-sensitive cortex correlated with attention and WM span
- Category activity coupled with control and memory regions for effective selection

## 1. Introduction

Working memory (WM) and cognitive control exhibit significant decline with normal aging (Salthouse, 2010), which can cause deleterious effects across cognitive domains (Park et al., 2002, 1996; Wingfield et al., 1988) and affect quality of life (Davis et al., 2010). These declines coincide with a predominant decrease in prefrontal cortex (PFC) volume (e.g. Haug and Eggers, 1991; Raz et al., 2005), which highlights the PFC as a prime candidate brain region that underlies the age-related changes in WM and cognitive control (Braver and Barch, 2002; West, 1996). Much research supports the idea that attention plays an important role in supporting WM function by guiding selection of relevant items amongst competing distractors during encoding and maintenance (Gazzaley and Nobre, 2012; Stokes and Nobre, 2012; Vogel et al., 2005; Vogel and Machizawa, 2004), but it is unclear whether age-related deficits in WM are owed to problems in selective attention.

Neuroimaging studies of WM function in aging have found age-related differences in the activation in PFC in older adults related to poor performance (e.g. Grady, 2008; Reuter-Lorenz et al., 2000; Rypma and D’Esposito, 2000), which might reflect problems with top-down control of attention in WM in aging (Hasher and Zacks, 1988; Zanto and Gazzaley, 2014). Some researchers propose that general age-related cognitive deficits, including problems with WM, can be attributed to a decline in inhibitory control and impairment in the inhibition of irrelevant information (e.g. Hasher and Zacks, 1988; Lustig et al., 2007; Zanto and Gazzaley, 2014). Older adults have poorer memory for target items than younger adults and better memory for irrelevant distractors (Rowe et al., 2006; also see Campbell et al., 2010), and exhibit stronger proactive interference in WM span tasks (Lustig et al., 2001; May et al., 1999). Relatedly, Braver (2012) suggested that age-related deficits are owed to declines in the active maintenance of task set for upcoming behavior, or ‘proactive control’ in older adults. Consequently, they rely more on ‘reactive control’, where the task set is retrieved only at the moment behavior is required, resulting in behavioral deficits in a variety of tasks. Finally, Gazzaley and colleagues showed that older adults had deficits in suppressing BOLD activity for irrelevant distractors in sensory cortex during selective WM tasks (Chadick et al., 2014; Gazzaley et al., 2005). Overall, these studies suggest that age-related deficits in WM could be a manifestation of poor top-down control for suppression of task-irrelevant information.

However, one line of behavioral work suggests that spatial attention is relatively preserved in aging (e.g. Greenwood et al., 1993; Nissen and Corkin, 1985; Tales et al., 2002), showing that older adults can use spatial pre-cues to improve perceptual performance as much as their younger counterparts. Recent studies also challenge absolute deficits in attention for WM in ageing, by showing that older adults are capable of orienting spatial attention within WM using retro-cues to improve WM performance (Mok et al., 2016; Souza, 2016) with relative preservation of the neural markers of this control (Mok et al., 2016). One study also suggests that feature-based retrospective attention is similarly preserved in aging (Gilchrist, Duarte, & Verhaeghen, 2016). In sum, it remains unclear whether WM deficits in aging are related to impairments in selective attention.

Here, we tested older adults on a selective WM task in the MRI scanner to examine older adults’ ability to use top-down, category-based attention to support WM performance, and to test whether neural markers of attention co-varied with selective WM performance. In addition, we administered a surprise subsequent recognition memory test to investigate the functional consequences of selective attention on memory for attended and ignored items. We recruited a relatively large sample of older adults for greater statistical power due to higher behavioral variability in older groups (Botwinick, 1978; Krauss, 1980; Welford, 1985), and to capitalize on this variability to test if the degree of top-down modulation during selective WM is correlated with behavioral measures of attention and WM, which might relate to mechanisms of successful cognitive aging. Examining variability within an older age group also allowed us to circumvent certain problems when comparing across age groups, such as motivation, fatigue, technology exposure, and cerebrovascular differences.

We found that older adults were able to use top-down attention to modulate category-sensitive cortex to support performance on a selective WM task. Selective attention to individual exemplars of face and house stimuli significantly affected subsequent recognition memory for these items, demonstrating that attention can have durable functional consequences in older adults. Category-sensitive cortex and the PFC were modulated according to task demands, and attention-related modulation was predictive of our behavioral measure of attention (effect of attention on subsequent memory) and with individual WM capacity within an older group.

## 2. Materials and Methods

### 2.1. Participants

Eighty-one healthy older adults (aged 60-87) were recruited from the community via local media and public advertisements. Of these, 75 participants completed the current experiment. Five participants were excluded from the analysis due to cortical abnormalities observed in their T1-structural scans. The remaining 70 participants (42 female) were 60-87 years old (68.5±0.85 years), had 16.0±0.44 years of education, scored ≥26 on the Mini-Mental State Examination (MMSE, Folstein et al., 1975), and had normal or corrected-to-normal vision and hearing. All participants were fluent in English. None of the participants had any current diagnosed psychiatric or neurological disorder, or were taking psychoactive medication. The study was approved by the Central University Research Ethics Committee of the University of Oxford and was carried out in accordance with the provisions of the World Medical Association Declaration of Helsinki.

### 2.2. Stimuli and Apparatus

Stimuli consisted of digital color photographs of faces and houses with 350×350 resolution (see task schematic in figure 1A and figure S1 for representative examples of the images used). Face stimuli were photographs of individuals of different ages (children, young, middle-aged, and elderly adults) and ethnicities. Only faces with neutral or positive expressions were selected. House stimuli were photographs of houses with a range of styles. The stimuli were chosen to be as natural and pleasant as possible to keep elderly participants engaged. There was no attempt to use standardized images, since this was not relevant to the aims of the study. All images were obtained via Creative Commons licence on Flickr. Seventy-two images were used in the main task (36 faces, 36 houses), and a separate set of 24 images (12 faces, 12 houses) was used in the practice session. In the subsequent memory test, 72 ‘old’ images that were presented in the fMRI scanner (36 faces, 36 houses) and 73 ‘new’ images that were not previously presented to participants (36 faces, 37 houses) were selected. Due to a technical error, one additional ‘new’ house image was presented (37 instead of 36).

The task was programmed and run in Presentation^®^ (version 16.2, www.neurobs.com). The task was presented using a display with spatial resolution of 1024 × 768 pixels and refresh rate of 60 Hz, projected onto a screen and reflected onto a mirror inside the scanner (horizontal length 26°).

### 2.3. Behavioral tasks

#### 2.3.1. Selective WM task

A selective-WM task with images of faces and houses tested the ability to focus on items from one category to guide WM performance (Figure 1A). In each block, participants selectively attended to the face or the house stimuli, and made a response (button press) when they saw an image from the attended category presented a second time within the same block (stimulus repetition). Images from the competing category were irrelevant and could be ignored.

At the start of each block, an instruction cue (2000 ms) indicated if the upcoming block required attention to faces (“REPEATED FACES”) or to houses (“REPEATED HOUSES”). Prior to each stimulus, a reminder (“F’ or “H”) was presented (2000 ms) to ensure participants were aware of the current task block, followed by the face or house stimulus (1000 ms). Each block consisted of ten images. For each stimulus, its associated target (stimulus repeat) could be presented after one to three intervening stimuli, or not presented at all. This was the case for both the attended and ignored stimulus categories, even though no response was required for the ignored category. Intervening stimuli were face or house images with equal probability. Stimuli from the attended blocks were not presented in the ignore blocks (e.g. faces in attend-face ignore-house blocks were not presented in the ignore-face attend-house blocks), and vice versa. Participants were not explicitly informed of these details but were simply asked to respond when they saw an image of the attended category presented a second time at any point during the block.

There were 18 trials for each stimulus-attention condition (attend-face, ignore-face, attend-house, ignore-house) with novel images (72 trials), and 12 trials for each condition with a repeated image (48 trials), giving 120 trials in total.

#### 2.3.2. Subsequent memory test

Participants were given a surprise subsequent-memory test after the main experiment, once they were outside of the scanner. On each trial, participants were presented with a face or house image which had either been shown in the main selective WM task (old) or an image they had not seen before (new). They judged whether they remembered seeing this image (old) or not (new) by making a forced-choice response on a keyboard. Participants were presented with 72 ‘old’ images (36 faces, 36 houses) and 73 ‘new’ images (36 faces, 37 houses) in a pseudo-randomized order. Half of the ‘old’ images were from the ‘attend’ blocks and the other half from the ‘ignore’ blocks. Images were the same across participants. One participant did not complete the memory test.

### 2.4. Experimental procedure

At the beginning of the session, the experimenter provided a demonstration of the task accompanied with verbal instructions from a prewritten script. Participants proceeded to perform the task in a practice session, which consisted of two attend-faces and two attend-houses blocks. The experimenter provided verbal feedback on their performance. Once the experimenter was satisfied that the participant understood the task, the participant proceeded with the task in the scanner.

For the main selective-WM experiment, participants completed six attend-faces blocks and six attend-houses blocks in the scanner in alternating order. In each block, there were ten trials, lasting a total of 30 seconds, followed by 14 seconds of rest with central fixation. Eye movements were not monitored. After the scan session, participants were given a subsequent memory test.

### 2.5. Behavioral data analysis

#### 2.5.1 Selective WM and subsequent memory

To characterise performance on the task in each block type, we computed the hit rate and mean correct reaction times for the face and house blocks separately. Hit rate was the number of responses when a target was presented (repeated stimulus in the attended category) divided by the total number of targets. To characterise the effect of category-based selective attention for WM on subsequent memory, we computed the proportion of correctly responded ‘old’ trials for stimuli that were presented in the fMRI session for attended faces, ignored faces, attended houses, and ignored houses separately. To examine subsequent-memory performance in general, we computed the proportion of correct responses for all stimuli in the memory test, separated into ‘old’ and ‘new’ stimuli. Participants correctly judged images as ‘old’ if they had been presented in the main experiment, and correctly judged images as ‘new’ if they were not presented previously.

To test the effect of category-based attention during selective WM on subsequent memory, a repeated-measures ANCOVA was performed on proportion ‘old’ stimuli remembered (responded ‘old’), with factors Attention (attend, ignore) and Stimulus Category (face, house), with covariates age, education, and gender. Follow up t-tests were used to test for differences between conditions, and Cohen’s d was used to determine effect sizes. Statistical analyses for behavioral performance were performed using Matlab R2015a, Matlab’s Statistics Toolbox, and R version 3.2.1 (R Core Team, 2015) using the afex package (Singmann et al., 2015).

#### 2.5.2. Neuropsychological Tests

Participants completed a neuropsychological battery and questionnaires separately to the current experiment. Individual differences in the ability to filter task-irrelevant distractors and enhance task-relevant items in WM tasks have been linked to differences in WM capacity (e.g. Linke et al., 2011; Vogel et al., 2005; Vogel and Machizawa, 2004). Therefore, we hypothesized that the ability to enhance processing of target stimuli and supress distractors during the selective-WM task would correlate with digit span, a standard measure of WM. Since there were three scores related to WM in the neuropsychology battery, we used an aggregate score by taking the mean of the forward, backward, and sequence spans. We tested for correlations between digit span and brain measures of attention using linear regression (in whole-brain fMRI analyses; see below) or Spearman’s partial correlation (rho), with age, education, and gender as covariates of no interest.

### 2.6. MRI acquisition

Functional and structural MRI data were acquired on a 3T Siemens TIM Trio System (Siemens, Erlangen, Germany) using a 32-channel head coil at the Oxford Centre for Clinical Magnetic Resonance Research (OCMR). An EPI-BOLD contrast image with 32 slices was acquired with 3-mm^3^ voxel size, repetition time (TR) = 2000 ms, and echo time (TE) = 30 ms. Flip angle was set to 78°. A fieldmap image was acquired with parameters as follows: 3.5-mm^3^ voxel size, TR = 488 ms, first TE = 5.19 ms, second TE = 7.65 ms. A high-resolution whole-brain T1-weighted structural image (MPRAGE) was acquired for registration purposes, with 1 mm^3^ voxel size, TR = 2040 ms, TE = 4.7 ms. A resting-state scan and a separate task scan were also acquired, which do not form part of this investigation.

### 2.7. fMRI analyses

fMRI data processing was carried out using FMRI Expert Analysis Tool (FEAT) Version 6.00, part of FSL (FMRIB Software Library; http://www.fmrib.ox.ac.uk/fsl). Pre-processing consisted of head-motion correction (MCFLIRT; Jenkinson et al., 2002), brain extraction (FSL’s Brain Extraction Tool; Smith, 2002), spatial smoothing using a Gaussian kernel of full width at half maximum (FWHM) 8 mm, and high-pass temporal filtering set at 275 seconds. Images were unwarped using B-0 fieldmaps (Jenkinson, 2004, 2003). Functional data were registered to standard space using FSL’s Boundary-Based Registration (BBR) and non-linear registration tools (Andersson et al., 2007; Greve and Fischl, 2009; Jenkinson et al., 2002; Jenkinson and Smith, 2001).

Time-series analysis was carried out using FMRIB’s Improved Linear Model (FILM) with local autocorrelation correction (Woolrich et al., 2001). Data were analysed using the general linear model (GLM). Ten explanatory variables were used to model task events: Face Attend Novel (FA; face stimulus in attend-faces block), Face Ignore Novel (FI; face in attend-houses block), House Attend Novel (HA; house in attend-houses block), House Ignore Novel (HI; house in attend-faces block), Face Attend Repeat (repeated face in attend-faces block), Face Ignore Repeat (repeated face in attend-houses block), House Attend Repeat (repeated house in attend-houses block), House Ignore Repeat (repeated house in attend-faces block), Instructions, Incorrect Trials. Stimulus EVs were 1000 ms with inter-stimulus intervals of 2000 ms. ‘Instructions’ modelled the initial instruction screen stating the block type (2000 ms). ‘Incorrect Trials’ included misses and false alarms (1000 ms). Since ‘Repeat’ conditions potentially required a response, they were modelled but not used in main analyses to avoid artefacts and confounds related to potential and actual responses. Time points affected by large head movements remaining after motion correction were identified by FSL’s Motion Outliers tool, and were included as confound regressors.

In the main analysis, four contrasts were used. To localise brain areas preferentially activated by face stimuli to identify region-of-interests (ROIs) in the fusiform gyri (FG), we tested for regions that evoked greater blood oxygen-level dependent (BOLD) activity for face compared house stimuli (FA & FI > HA & HI). To localise areas preferentially active for house stimuli to identify ROIs in the parahippocampal gyri (PHG), we tested for regions that evoked greater activity for house stimuli compared face stimuli (HA & HI > FA & FI; see Region-of-interest analyses section for details). To test for areas that were modulated by attention during selective WM for faces, we contrasted Face Attend with Face Ignore (FA > FI). For selective WM for house stimuli, we contrasted House Attend with House Ignore (HA > HI). Note that the contrasts used to define face-sensitive and house-sensitive areas were independent from the attention contrasts and therefore do not suffer from selection bias (Kriegeskorte et al., 2009).

Group-level analyses were carried out using FMRIB’s Local Analysis of Mixed Effects (FLAME; Woolrich et al., 2004). Z (Gaussianized T/F) statistic images were thresholded using clusters determined by Z>2.3 and a whole-brain corrected, family-wise error (FWE) cluster significance threshold of p=0.05 (Worsley, 2001). Grey-matter images were extracted from each participant’s structural scan using FMRIB’s Automated Segmentation Tool (FAST; Zhang et al., 2001) and added as a covariate to account for voxel-wise differences in grey matter in all group analyses.

### 2.8. Conjunction analysis

A conjunction masking analysis on the group-level z-statistic images (FWE cluster-corrected) was performed to localize attentional control areas involved in both face attention (FA>FI) and house attention (HA>HI). To compute the conjunction image, we multiplied the FA>FI contrast z-statistic image with the HA>HI contrast z-statistic image. Each contributing contrast was thresholded to include only significant clusters at the whole-brain level, so that voxels that do not overlap in the two images would have a value of zero. The resulting image was binarized. Only voxels that were significant at Z > 3.09, or p < 0.001 in both contrasts were selected.

### 2.9. Region-of-interest analyses

To test for attentional modulation in category-sensitive areas, we focused on the posterior fusiform (FG; Allison et al., 1994; Kanwisher et al., 1997; Puce et al., 1995) within the network of areas that respond strongly to faces (Allison et al., 1999; Behrmann and Plaut, 2013). To test for modulation in place-sensitive areas, we focused on the parahippocampal gyri (PHG; Epstein and Kanwisher, 1998). To do this, the coordinates of the peak BOLD signal (z-statistic) at the group-level were identified in the inferior temporal (IT) lobe from the faces versus houses contrast (FG) and the houses versus faces contrast (PHG). This gave us one ROI in each hemisphere for each contrast, giving four ROIs per participant. Masks centred on the peak in each ROI in standard MNI space (17 mm^3^ cubes) were transformed into single-subject space. We ensured no spatial overlap between the masks in all participants, and that each participant’s ROIs laid within the vicinity of the face and place-sensitive areas (Epstein and Kanwisher, 1998; Kanwisher et al., 1997). Within each mask, we extracted the coordinates with the peak activity (z-statistic) from each participant for each localiser contrast. A pair of spherical masks (5 mm^3^) were constructed and centred on the single-participant defined FGs and PHGs from each hemisphere to extract mean beta estimates in the FA, FI, HA, and HI conditions. Since we were mainly interested the effect of attention on BOLD activity in category-sensitive cortex, and not hemispheric differences, beta estimates were averaged across hemispheres in the FG and the PHG in each condition.

To examine top-down modulation during category-based attention in face-sensitive and house-sensitive regions, repeated-measures ANCOVAs were performed on BOLD activity (beta estimates) for each bilateral ROI (FG, PHG) with factors Attention (Attend, Ignore) and Stimulus Category (Faces, Houses); with age, education, and gender as covariates. Follow-up pairwise t-tests were used to assess condition differences. We tested for a relation between attentional modulation (BOLD in attend minus ignore conditions in the category-sensitive bilateral ROIs) and the effect of attention on subsequent memory (memory accuracy difference between attended images versus ignored images) using Spearman’s partial correlation, controlling for age, education, and gender. We also tested for a relation between attentional modulation and digit span, controlling for age, education, and gender. Finally, we also tested if attentional modulation was correlated with age, controlling for education and gender. Correlation coefficients were compared by using Fisher’s r-to-z transformation and tested for significant differences (Cohen and Cohen, 1983). Participants were excluded from a certain statistical test if the extracted BOLD activity in a given condition, or from a contrast of two attention conditions, was greater or smaller than three standard deviations from the group mean. Using this criterion, we excluded five participants in total: one participant with an outlier in the FG in condition FI, one participant from the FG in the FA-FI contrast, one participant from the PHG in condition HA, one participant in the FG in condition FI, and one participant in the PHG in condition FA and the FA-FI contrast (same participant). Results were equivalent with or without outlier exclusion.

### 2.10. Brain-behavior correlations within attention-related modulated regions

To test for individual differences in attention-related modulation of BOLD for faces (FA>FI) and houses (HA>HI) that corresponded to i) the effect of attention on subsequent memory, ii) WM capacity, and iii) age; subsequent-memory difference scores for faces (proportion remembered attended face images – proportion remembered ignored face images) and houses (proportion remembered attended house images – proportion remembered ignored house images), mean digit span, and age were included as regressors in the GLM. Education and gender were included as regressors of no interest. To test for relations between these measures and activity in attention-modulated regions, we ran a group analysis with only education and gender as covariates of no interest and constructed masks from two contrasts (FA>FI and HA>HI, separately) to localize the areas that showed significant attentional modulation. Two group-level analyses for each contrast were performed with age as a regressor and education and gender as covariates of no interest, using the activation masks as pre-threshold masks to restrict the correlations to areas that exhibited attentional modulation in each respective contrast. To test the relation between areas that showed significant attentional modulation with digit span and subsequent memory, a group analysis was conducted with age, education, and gender as covariates to obtain the activation masks for the two main contrasts. Group-level analyses were then performed to test for a relation between the attentional modulation for faces (FA>FI) and digit span using the activation mask from the face attention contrast, and attentional modulation for houses (HA>HI) and digit span using the activation mask from the house attention contrast. The activation mask from the face-attention contrast was used to test for a relation between the attentional modulation for faces and the subsequent memory difference score for faces, and the activation mask from the house-attention contrast was used to test for a relation between modulation for houses and the subsequent memory difference score for houses. In both analyses, age, education, and gender were included as covariates of no interest. A group-level analysis without pre-threshold masking produced very similar results.

Information on significant clusters including peak locations, cluster size, statistical values, and peak MNI coordinates is reported in the Appendix (fMRI cluster tables). The first local maximum value reported for each cluster is the peak activation within that cluster. The Harvard-Oxford cortical and subcortical structural atlases (in FSL) were used as a guide for labelling anatomical locations of the local maxima.

### 2.11. Psychophysiological interaction analyses

We used psychophysiological interaction (PPI) analyses (Friston et al., 1997) to test for brain areas that were functionally coupled with category-sensitive visual cortex during selective WM. PPI was implemented in FSL (O’Reilly et al., 2012).

We tested for areas that showed greater coupling with bilateral FG (as PPI seed) when encoding faces that were attended versus ignored (FA > FI). We performed the same analysis for bilateral PHG for attended versus ignored house stimuli (HA > HI). For each PPI, we included four regressors: 1) the contrast of interest (e.g. FA – FI); 2) the time course of the seed ROI (e.g. bilateral FG); 3) the element-wise product of the first two regressors, representing the PPI; and 4) both conditions (e.g. FA + FI). The fourth regressor was included to model the shared variance of the two regressors not modelled by the contrast itself. In each analysis, all other task regressors noted in the main analysis were also included (O’Reilly et al., 2012). Group-level analyses were performed using FLAME, with age and digit span as regressors of interest, and education and gender as regressors of no interest. Group-level analyses were performed separately with subsequent memory difference scores for faces as a regressor for the FA>FI contrast (PPI with FG), and subsequent memory difference scores for houses as a regressor for the HA>HI contrast (PPI with PHG). Since we were interested in brain regions that correlated with these measures within and outside of areas that showed attentional modulation, pre-threshold activation masking was not used.

## 3. Results

### 3.1. Behavioral results

#### 3.1.1. Selective WM task performance

Performance accuracy was near ceiling for both attend-face (hit rate: 0.97±0.01) and attend-house (hit rate: 0.95±0.01) blocks (hit-rate difference: t(69)=1.80, p=0.08, d=0.25), and participants were slightly faster during attend-face (628±8.93 ms) compared to attend-house blocks (650±9.96 ms; t(69)=-2.05, p=0.045, d=-0.28). Performance was not significantly correlated with age in the attend-face blocks (hit rate: rho=-0.07, p=0.60; RT: rho=0.02, p=0.88), attend-house blocks (hit rate: rho=0.10, p=0.41; RT: rho=-0.07, p=0.55), or the difference between attend-face and attend-house blocks (hit rate: rho=-0.15, p=0.21; RT: rho=0.06, p=0.64).

**Figure 1.**
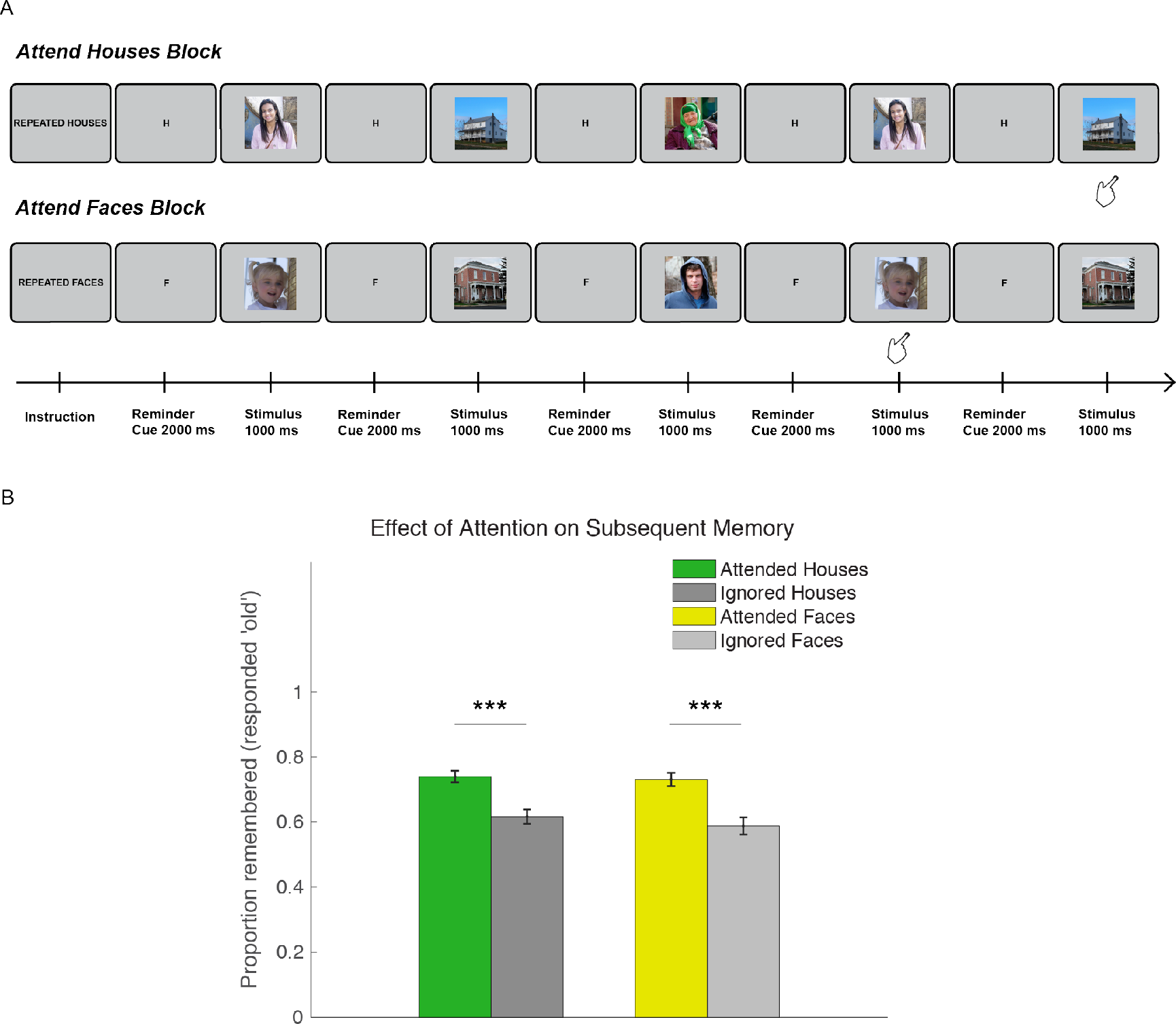
Task schematic and behavioral results on subsequent-memory test. (A) A selective WM task was performed in the fMRI scanner. In each block, participants monitored a sequence of intermixed face and house stimuli, and attended to one of the stimulus categories. In the attend-houses blocks (top), participants made a response when a house stimulus appeared a second time, ignoring the faces. In the attend-faces blocks (bottom), participants responded when a face stimulus appeared a second time, ignoring the houses. A cue “H” or “F” preceded each stimulus to remind the participant of the current task block. These particular stimulus sequences are for illustration only, and were different to those used in the experiment. (B-C) Subsequent memory performance. (B) Bar plot showing proportion correct (responded ‘old’) for attended houses presented during the attend-house blocks (green), ignored houses during the attend-face blocks (dark grey), attended faces presented during the attend-face blocks (yellow), and ignored faces presented during the attend-house blocks (light grey). Note that attended and ignored stimuli are all ‘old’. Error bars represent standard error of the mean. Asterisks denote significant pairwise differences. *** p<1e-06.

#### 3.1.2. Subsequent memory performance

Older adults performed relatively well on the subsequent memory test for faces (proportion correct for attended and ignored old images and new images: 0.78±0.01) and houses (0.71±0.01). They were much more likely to remember images that were attended (0.74±0.02) in the main experiment compared to images that were ignored (0.60±0.02) (F(1,65)=57.78, p=1.49e-10, η_p_^2^=0.47; Figure 1B), reflecting effective selection of relevant items and suppression of distractors during the WM task. There was no significant difference for memory between faces and houses (F(1,65)=0.90, p=0.35, η_p_^2^=0.01) and no difference in the attention effect between stimulus categories (Interaction: F(1,65)=0.62, p=0.43, η_p_^2^=0.009).

### 3.2. fMRI Results

#### 3.2.1. Common activations in category-based selective WM

Older adults recruited a common set of brain regions during selective WM for faces and during selective WM for houses, including PFC, temporal-occipital cortex including IT gyrus, fusiform gyrus, and lateral occipital cortex (LOC) (Figure 2). Areas activated in both hemispheres included frontal operculum (fO) and the insular cortex, frontal pole, dorsal anterior cingulate cortex (ACC), FG, IT gyrus, and LOC. Activations in the left hemisphere included middle frontal gyrus (MFG), precentral gyrus, inferior frontal gyrus (IFG) par opercularis, and orbital frontal cortex (OFC). Common activations in subcortical areas included small regions in the right caudate, left hippocampus, thalamus, and amygdala (not shown in figure).

**Figure 2.**
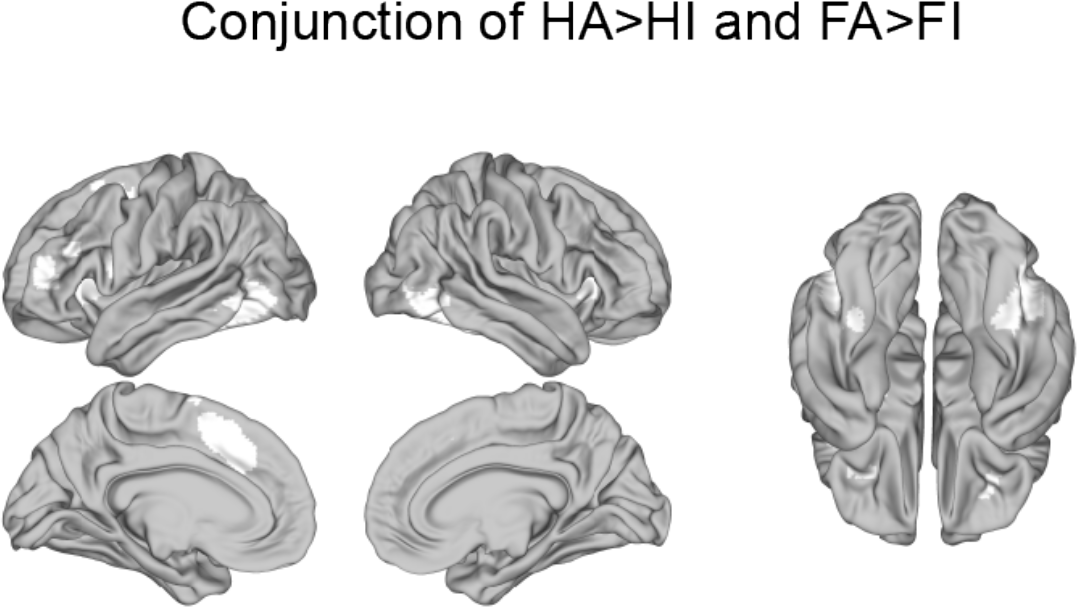
Common brain regions recruited during category-based selective WM for both faces and houses in older adults. Conjunction mask (white) showing recruitment of dorsolateral and inferior prefrontal cortex and lateral occipital cortex during selective WM. Areas in white mark regions that were active for both face attention (FA>FI) and house attention (HA>HI) (Z > 3.09 voxel-wise uncorrected within significant clusters in both contrasts). Brains are displayed in neurological convention; L=L.

#### 3.2.2. Top-down modulation of category-sensitive visual cortex during selective WM in aging

Older adults were able to use top-down attention to modulate category-sensitive visual cortex during selective WM (Figure 3A). Replicating the well established pattern of categorical functional specialization, BOLD activity was stronger for faces in the FG (F(1,63)=304.98, p=7.86e-26, η_p_^2^=0.83) and houses in the PHG (F(1,64)=461.87, p=5.68e-31, η_p_^2^=0.88). Both regions were strongly modulated by attention to stimulus category. There was a significant effect of category-based attention in the FG (F(1,63)=34.99, p=1.48e-07, η_p_^2^= 0.36), with greater BOLD for attended versus ignored stimuli, but no significant interaction with Stimulus Category (F(1,63)=2.74, p=0.1, η_p_^2^=0.04). There was a significant effect of attention for houses in the PHG (Attention: F(1,64)=20.16, p=3.03e-05, n_p_^2^=0.24) and an interaction between Attention and Stimulus Category (F(1,64)=19.60, p=3.80e-05, η_p_^2^=0.23) where BOLD activity was greater for the attended-houses compared to ignore-houses conditions (t(69)=6.37, p=1.91e-08, d=0.52) but not in the attend-faces versus ignore-faces conditions (t(69)=1.37, p=0.18, d=0.13; difference between conditions: t(67)=4.48, p=2.88e-05, d=0.62).

The degree of top-down modulation in bilateral FG positively correlated with effect of attention on subsequent memory for faces (memory difference between attended versus ignore face images; rho=0.37, p=0.003; Figure 3B, right). Given the lack of an interaction of Attention and Stimulus Category in bilateral FG in the ANCOVA above, we might expect top-down modulation in FG to correlate with the effect of attention on house memory as well. Alternatively, if the modulation in FG was more specific to encoding faces into memory, this should only correlate with attention effects on memory for faces. We found that top-down modulation for houses in bilateral FG was not significantly correlated with attention effects on memory for houses (rho=0.02, p=0.86) and the correlation coefficient was significantly greater for the effect of attention of face memory compared to house memory (z=2.08, p=0.038). Top-down modulation in bilateral PHG positively correlated with digit span (rho=0.34, p=0.005; Figure 3C, right). Top-down modulation in bilateral PHG was not significantly correlated with subsequent memory for houses (rho=0.17, p=0.17). Age was not significantly associated with top-down modulation in FG (rho=-0.05, p=0.67) or in PHG (rho=-0.01, p=0.91)

Among brain regions that showed attention-related modulation during selective WM for faces, we found that activity in right posterior IT including the LOC and posterior FG positively correlated with the effect of attention on subsequent memory for faces (p=0.001, cluster-corrected; Figure 3B, left; Table S1). Among attention-modulated regions during selective WM for houses, activity in right LOC and bilateral IT including the PHG was positively correlated with digit span (p’s<0.026, cluster-corrected; Figure 3C, left; Table S2). Neither showed a significant relationship with age (all p>0.05, cluster-corrected). The regions that showed significant positive correlations with digit span and subsequent memory difference scores were qualitatively very similar when analyses were performed with or without a pre-threshold mask, with correlations mainly restricted to visual areas.

**Figure 3.**
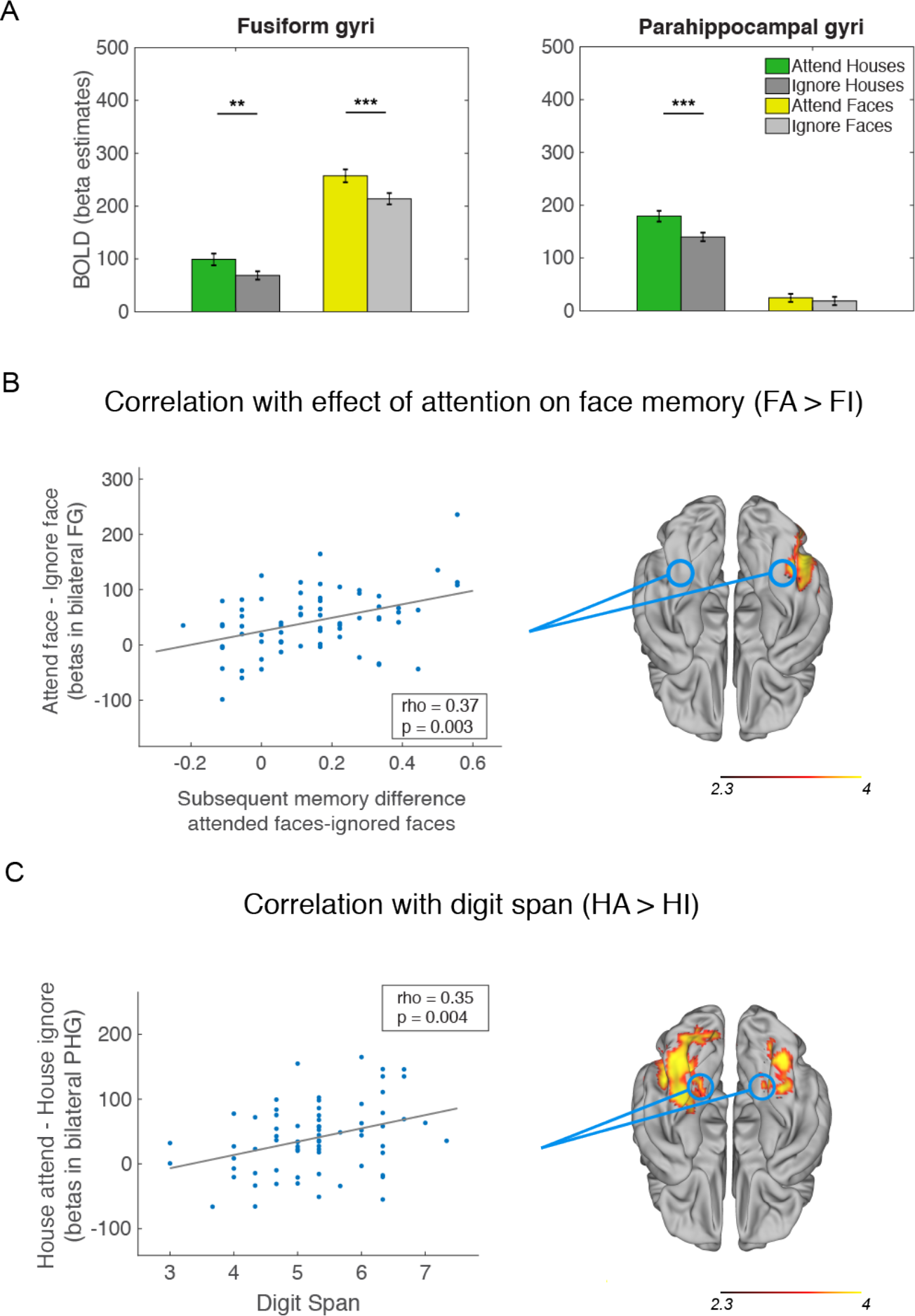
Top-down attentional modulation in category-sensitive visual cortex during selective WM and correlations with behavior. (A) Bar plots showing attention-related modulation of category-sensitive visual cortex (ROI analyses). Left: BOLD activity in bilateral fusiform gyri (FG) was modulated by stimulus category, with greater activity for faces than houses, but attention-related modulation was similar for faces and houses. Bilateral FG showed greater activity in the attend-face (FA; yellow) versus the ignore-face (FI; light grey) conditions, as well as in the attend-house (HA; green) compared to the ignore-house (HI; grey) conditions. Right: BOLD activity in bilateral parahippocampal gyri (PHG) was modulated by stimulus category, with greater activity for houses than faces, and displayed strongly selective attention-related modulation for WM encoding of houses. Bilateral PHG showed greater activity in the attend-house (HA; green) compared to the ignore-house (HI; grey) conditions, but not for attend-face (FA; yellow) versus the ignore-face (FI; light grey) conditions. ***p<7.89e-08; **p=0.0003. Beta estimates are in arbitrary units. Error bars represent standard error of the mean. (B-C) The degree of attention-related modulation was positively correlated with the effect of attention on subsequent memory and with digit span. B) Left: Scatterplot of BOLD activity (FA-FI) from participant-defined bilateral FG plotted as a function of subsequent memory difference scores for faces. Right: Degree of attention-related modulation in right lateral occipital cortex (LOC) and FG (approximate locations circled) during selective WM for faces was positively correlated with the effect of attention on subsequent memory for faces. Significant regions are displayed in red-yellow shading (p=0.001, cluster-corrected). C) Left: Scatterplot of BOLD activity (HA-HI) from participant-defined bilateral PHG plotted as a function of digit span. Right: Degree of attention-related modulation in bilateral IT, including PHG (approximate locations circled) and left LOC during selective WM for houses was positively correlated with digit span, a measure of WM capacity obtained in a separate session. Significant regions are displayed in red-yellow shading (p’s<0.026, cluster-corrected). Activations on brains are z-statistic images. Brains are displayed in neurological convention; L=L.

#### 3.2.3. Functional coupling with category-sensitive visual areas during selective WM

To identify control-related brain regions in which activity co-varied with activations in category-sensitive cortex depending on category-based selective attention, we used a PPI analysis. Activity in bilateral FG was coupled with bilateral precuneus and cuneal cortex, left superior parietal lobe, and right postcentral gyrus when participants were ignoring faces compared to attending faces (FI > FA; p’s < 0.02, cluster-corrected; Figure 4B; Table S3). Although no brain areas showed significant coupling with bilateral FG for face attention (FA>FI) at the group level, the degree of this coupling in a network of brain areas was positively correlated with the effect of attention on subsequent memory for faces (subsequent memory difference score), including the right hippocampus, LOC, superior temporal sulcus, OFC (figure 4C, p’s < 0.05, cluster-corrected; Table S4). In other words, participants that showed a larger difference between subsequent memory for attended faces compared to ignore faces showed more coupling between bilateral FG and these brain areas during selective encoding of faces into WM. Activity in bilateral PHG preferentially coupled with the left PFC, including the fO, IFG pars opercularis, central opercular cortex, and precentral gyrus during attending relative to ignoring houses (Figure 4A; p=5.36e-7, cluster-corrected; Table S5).

**Figure 4.**
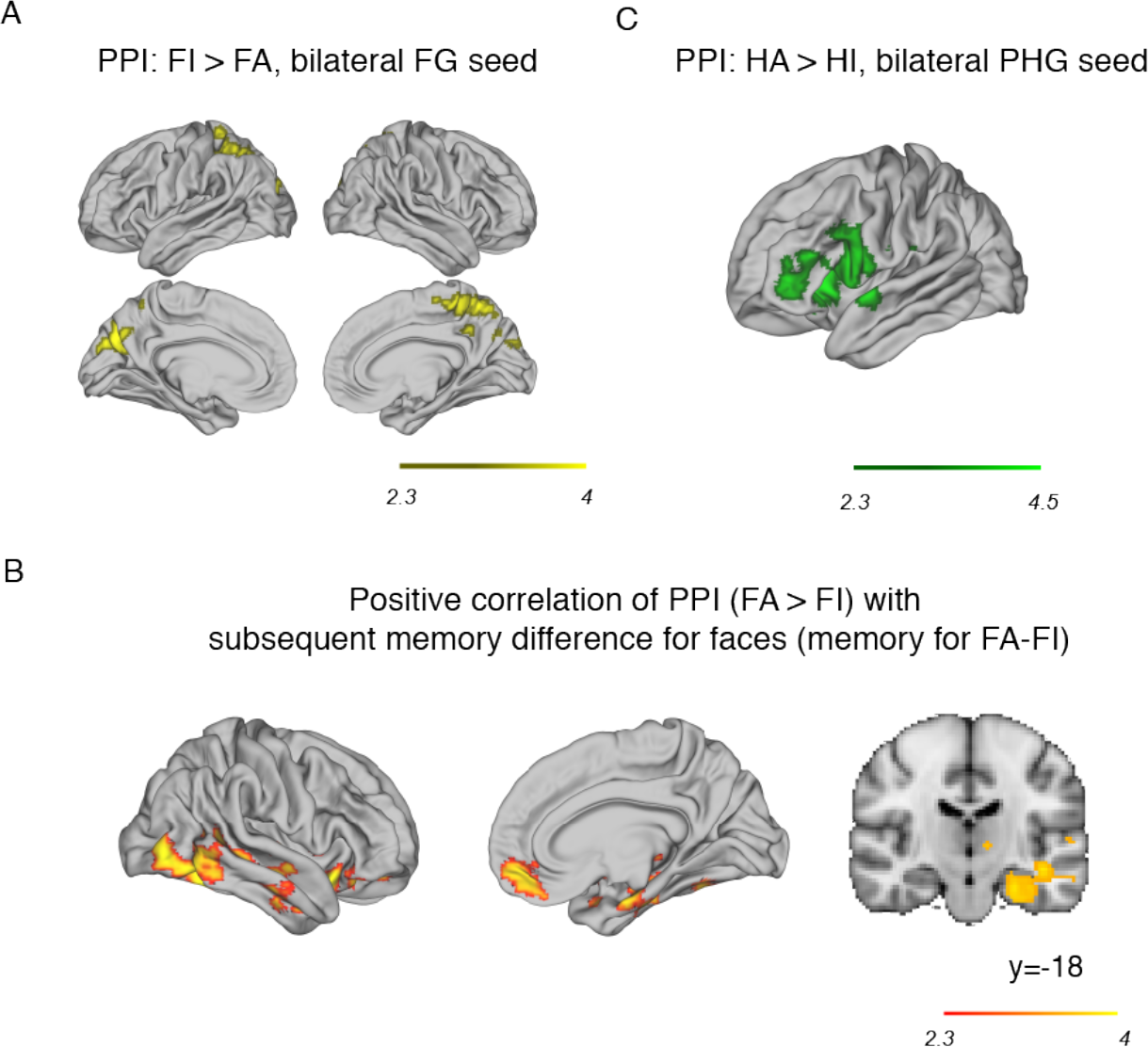
Psychophysiological interaction analyses showing brain areas that exhibited coupled activity with category-sensitive visual areas during selective WM in older adults. A) Activity in bilateral precuneus and left superior parietal lobe were more coupled with bilateral FG during suppression of distractor faces relative to encoding faces into WM (FI>FA, p=1.19e-7, cluster-corrected). B) The degree of coupling between bilateral FG and a network of frontal and temporal brain regions predominantly on the right hemisphere during selective WM for faces (FA>FI) were positively correlated with the effect of attention on subsequent memory for faces (FA-FI; red-yellow shading, p’s < 0.05 cluster-corrected). y-coordinate in MNI space. C) Activity in left fO/anterior insular, IFG, and precentral gyrus, was coupled with activity in bilateral PHG during selective WM encoding of house stimuli (HA>HI; p=5.36e-7, cluster-corrected). Brains are displayed in neurological convention; L=L.

## 4. Discussion

We tested a large group of older adults and found that they were able to use top-down control to modulate category-sensitive cortex in a selective WM task. Older adults used category-based attention to encode target items into WM and suppress distractor items. Their effective use of selective attention was evident from their high degree of accuracy during task performance. Better subsequent memory of attended images relative to ignored images further showed that selective attention to individual items had functional consequences for subsequent recognition memory. Category-sensitive cortex showed greater activation to the attended versus ignored category during WM encoding, and the degree of modulation was associated with behavioral consequences of selective attention and WM span. Purported control regions, including dorsolateral PFC, IFC, and ACC, were recruited in both types of category attention in concert with category-sensitive cortex, and the degree of coupling between FG and the right hippocampus, STS, and OFC was correlated with the effect of attention on subsequent recognition memory for faces.

As a group, older adults recruited greater activity for attended compared to ignored faces in the FG and attended compared to ignored houses in the PHG, as reported in other studies in younger (O’Craven et al., 1999) and older adults (Chadick et al., 2014; Gazzaley et al., 2005). Attention-related modulation in bilateral PHG was reliably category-selective, with greater activity in attend-compared to ignore-house conditions but not for attend-compared to ignore-face conditions. Modulation in the FG was less selective, with a similar degree of modulation irrespective of stimulus category, despite overall significantly lower activity for house stimuli. Interestingly, the effect of attention on subsequent memory for faces (memory for attended minus ignore faces) was only correlated with modulation in the FG but not the PHG,indicating some attention-related category specialisation after all. In the PHG, there was a strong positive correlation between attention-related modulation and WM span as measured by digit span tasks. Since WM span was assessed using a different task and in a separate session, this suggests that attention-related modulation in category-sensitive cortex can also index general WM function. This extends previous work that showed a positive correlation between attention-related suppression in PHG with WM performance in the same task (Gazzaley et al., 2005; see below for further discussion), and gives further support to the idea that selective attention plays an important role in supporting effective WM function (Myers et al., 2017; Nobre and Stokes, 2011; Vogel et al., 2005; Vogel and Machizawa, 2004) in older adults.

The PFC plays a key role in top-down attention (Miller and Cohen, 2001; Petersen and Posner, 2012), and inferior PFC regions such as the IFG appear to play a key role in category and feature-based attention in younger adults (Baldauf and Desimone, 2014; Zanto et al., 2011, 2010). Consistent with these studies, we found strong recruitment of activity in PFC during selective WM for faces and for houses in older adults, including bilateral IFC (including IFG), left dorsolateral PFC, and ACC. During selective WM for houses, bilateral PHG showed coupled activity with the left IFC, similar to studies that find IFG involvement in category and feature-based attention (Baldauf and Desimone, 2014) and selective WM tasks (Zanto et al., 2010, 2011). We also found that during face attention, activity in the FG for suppression of faces (FI>FA) was coupled with greater activity in task-negative areas including the precuneus (Chadick and Gazzaley, 2011; Kim et al., 2010). Areas implicated in explicit memory encoding were coupled with face-sensitive FG during face attention and correlated with subsequent memory advantages. Specifically, the degree of coupling between the right hippocampus, STS, and OFC with FG was predictive of memory for relevant (attended) faces over distractors, which suggests that effective top-down attention can modulate long-term memory encoding in older adults (see Aly and Turk-Browne, 2018). Notably, there was no hint of a relation between this effect with age in our older sample, meaning this was not simply a younger subset of the older adults who were performing better. The hippocampus is involved in long-term memory (Nadel and Moscovitch, 1997) and the right hippocampus has been implicated in face memory in younger adults (Grady et al., 1995; Haxby et al., 1996; Milner, 1968). Grady et al. (1995) showed activation of the right hippocampus for encoding faces into memory in younger but not in older adults, which they suggested could have been the reason for poorer face memory in the older group. In light of this, our results suggest that the cognitively healthy adults within our current group may have been able to recruit this memory network more effectively for selective encoding of faces into memory. Our findings suggest that, if older adults are able to recruit brain regions effectively and in concert with each other, they can more effectively encode information into WM amongst distractors, which may reflect healthier or more ‘youth-like’ cognitive brain processes.

Some researchers have attributed general age-related cognitive deficits specifically to a decline in inhibitory control (Hasher and Zacks, 1988; Lustig et al., 2007), and impairment in the inhibition of irrelevant information (e.g. Hasher and Zacks, 1988; Zanto & Gazzaley, 2014; also see McNab et al., 2015). Gazzaley et al. (2005) tested older and younger adults on a selective WM task with face and scene images using fMRI and showed that the older group did not suppress BOLD activity according to attentional demands. Specifically, they concentrated on the left PHG, and found that brain activity was supressed when scenes were distractors (relative to passively viewed scenes) in younger but not in older adults, whereas left PHG activity was enhanced during encoding scenes into WM (compared to passively viewed scenes) in both groups (also see Chadick et al., 2014). In one study Gazzaley et al. (2005) found a relation between the degree of suppression in the left PHG with WM accuracy for faces, and in a second study (Chadick et al., 2014) they found that suppression in the left PHG was negatively correlated with distractibility (face WM accuracy with distractors minus face WM accuracy without distractors) in relatively small samples of older adults (18 and 16, respectively). It appears that enhancement effects were not related to behavior (not reported), and notably, enhancement effects were not significantly different between age groups. In our group of older adults, we found strong bilateral attention-related modulations during selective WM for both faces and houses, and a significant correlation between modulation and behavioral markers of attention and WM. Our strong effects are likely due to a large cohort which could have overcome some problems such as lower reliability related to MRI signals, and higher behavioral variability in older adults groups (Botwinick, 1978; Krauss, 1980; Welford, 1985). This suggests that it is possible to examine brain markers of effective top-down control for selective WM in older adults using our experimental design, which is an elderly-friendly task and more efficient (requires fewer experimental conditions), and allows testing large groups of participants to more effectively probe inter-individual differences.

In the current study, older adults showed strong recruitment of non-sensory brain regions implicated in cognitive control and memory, along with appropriate modulation in category-sensitive cortex which positively correlated with behavioral performance but not with age. Some studies have reported older adults show strong or over-recruitment of control areas including frontal and parietal cortex (e.g. Cabeza et al., 2002; Cappell et al., 2010; Madden, 2007), whereas others reported under-recruitment of control areas (e.g. Nyberg et al., 2010; Rypma and D’Esposito, 2000). Reuter-Lorenz and Cappell (2008) suggested these results might reflect the ability of older adults to cope with the task demands. Specifically, when older adults are proficient at a particular task, there may be over-recruitment of PFC, reflecting the greater cognitive resources required to perform to a similar level to younger adults, whereas when they struggle, they would show under-recruitment reflecting poor task execution (Cappell et al., 2010; Schneider-Garces et al., 2010). Indeed, mounting evidence suggests that the degree of activation of control regions is linked more closely with performance than age (e.g. Cappell et al., 2010; Höller-Wallscheid et al., 2017; Kurth et al., 2016; Loosli et al., 2016; Nagel et al., 2011, 2009). Our results are consistent with this work, suggesting that the degree of recruitment of purported control regions and task-relevant attention-related modulation play a role in supporting cognitive performance, and may be a marker of the cognitive health of these brain mechanisms in normal aging.

Our study shows that the ability to use selective attention to control the contents of WM is relatively preserved in aging and could be used to support other cognitive processes that undergo age-related decline. Despite effective selective WM in older adults as a group, there were individual differences in the degree of top-down modulation in relevant brain regions, and this variability was predictive of WM function. Our study suggests that studying the variability in behavior and brain patterns in older adults is a promising way to characterise preserved versus impaired cognitive abilities in aging, and future studies should use multiple tasks with varying levels of cognitive demand to reveal how the brain markers adapt or break down in different cognitive scenarios in normal aging.

## Acknowledgements

We thank Susie Murphy, Angela Rylands, and Emily Holmes for their help on setting up the Cognitive Health in Ageing (CHA) project, and Claire Burley and Clare Palmer for help with recruitment. This study was supported by a Wellcome Trust Senior Investigator Award (ACN) 104571/Z/14/Z, a European Union FP7 Marie Curie ITN Grant N. 606901 (INDIREA), the National Institute of Health Research (NIHR) Oxford Health Biomedical Research Centre (BRC) based at the Oxford Health Foundation Trust and the University of Oxford. The Wellcome Centre for Integrative Neuroimaging is supported by core funding from the Wellcome Trust (203139/Z/16/Z). The views expressed here are those of the authors and not necessarily those of the NHS, the NIHR or Department of Health.

## Appendix

**Figure S1.**
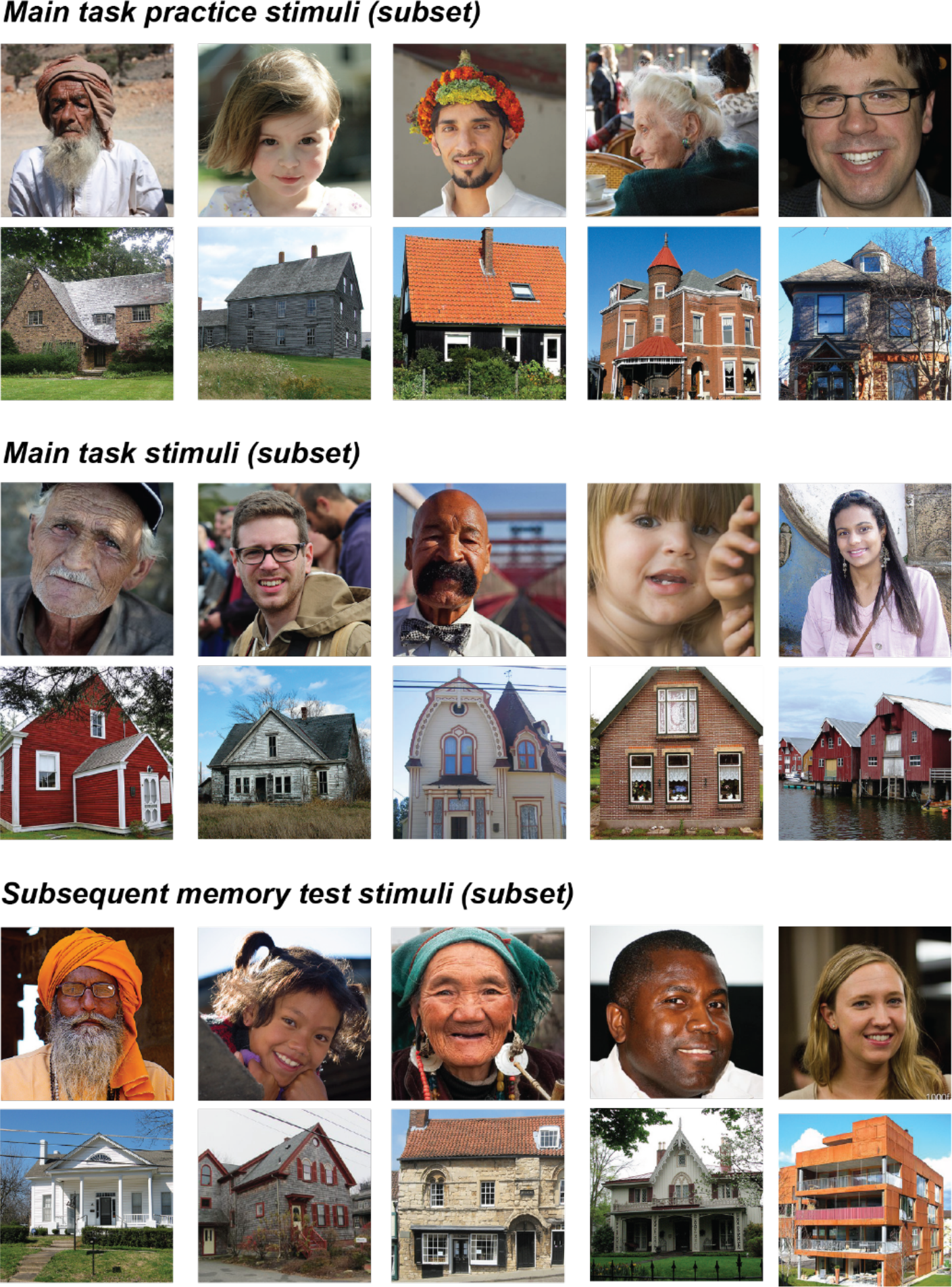
Examples of stimuli used in the tasks. Top: Subset of the stimuli used in the practice session of main experiment (selective WM task). Middle: Subset of the stimuli used in the main experiment. Subset of the stimuli (‘new’ images not presented in the main experiment) used in the subsequent memory test.

### fMRI cluster tables

Naming guided by the Harvard-Oxford cortical and subcortical structural atlases

#### Abbreviations

fO: : frontal operculum
IFG: : inferior frontal gyrus
ITG: : inferior temporal gyrus
LO: : lateral occipital cortex
MTG: : middle temporal gyrus
OFG: : occipital fusiform gyrus
SPL: ; superior parietal lobule

**Table S1.** Peak anatomical locations, Z values, and MNI coordinates from the significant cluster which exhibited a significant correlation between attentional modulation in the right LO and IT cortex for faces (FA > FI) and the subsequent memory difference score for face images (within brain regions that showed significant attentional modulation for FA > FI; with activation mask from the FA > FI contrast; see Methods).

| Location | Cluster number | Z value (local maxima) | x | y | z |
| --- | --- | --- | --- | --- | --- |
| Right ITG | 1 (cluster extent: 1066 voxels, $p = 0.001$) | 3.83 | 50 | -54 | -12 |
| Right inferior LO | 1 | 3.25 | 50 | -68 | -10 |
| Right ITG | 1 | 3.18 | 44 | -36 | -14 |
| Right ITG | 1 | 3.13 | 58 | -38 | -14 |
| Right inferior LO | 1 | 2.99 | 46 | -78 | 2 |
| Right inferior LO, MTG | 1 | 2.92 | 44 | -60 | 8 |

**Table S2.**
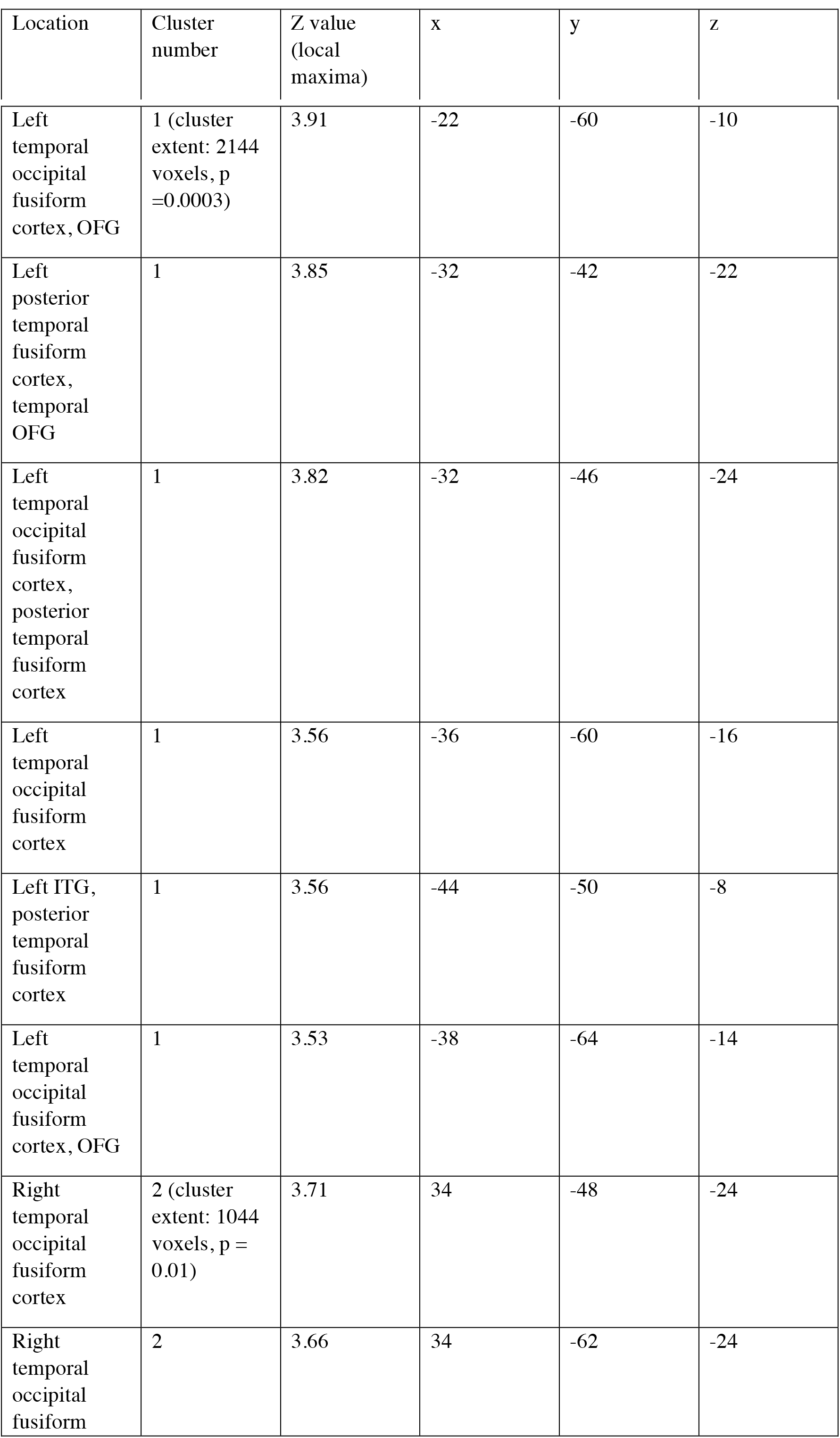
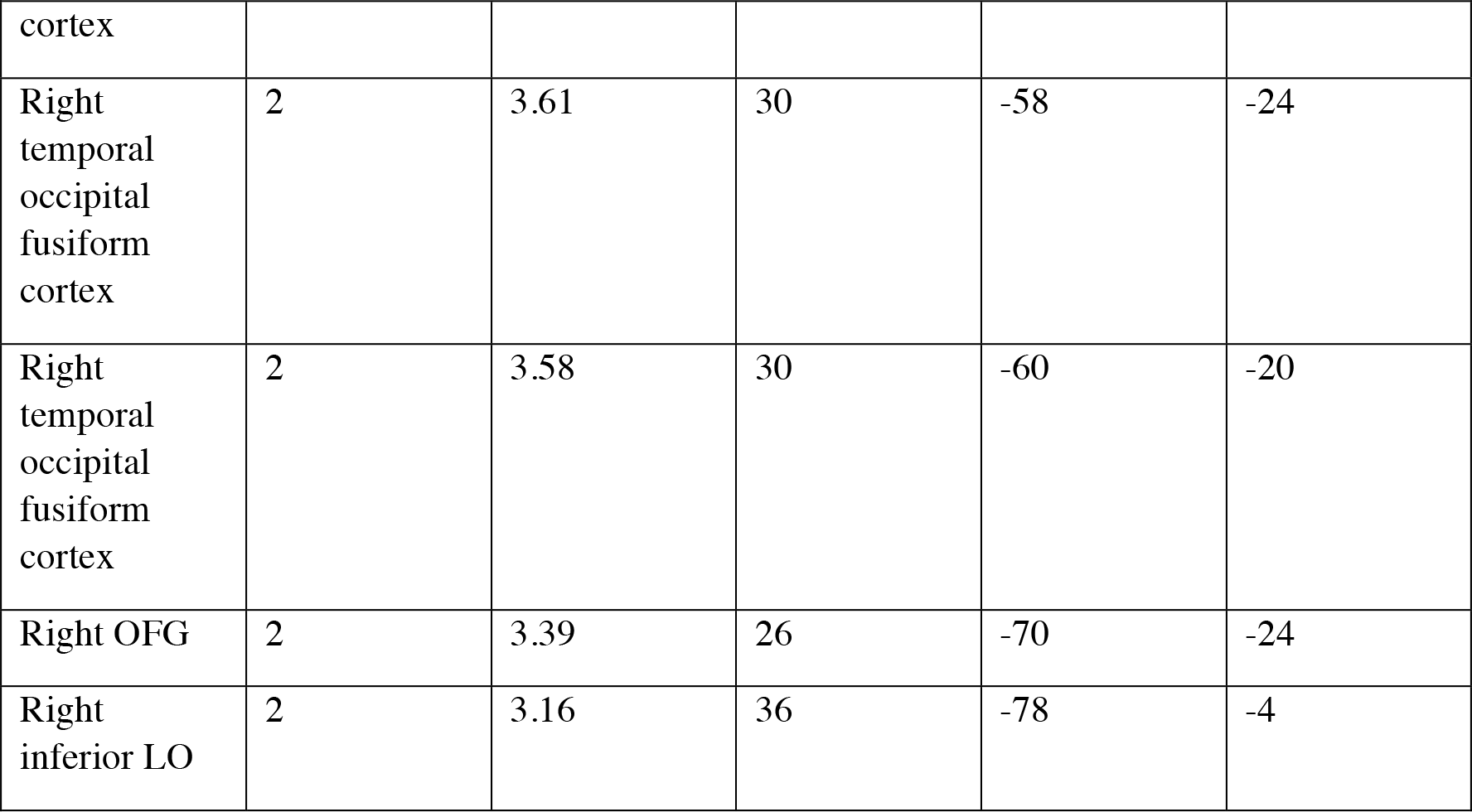
Peak anatomical locations, Z values, and MNI coordinates from the significant clusters which exhibited a significant correlation between attentional modulation in the PHG and IT cortex (HA > HI) and digit span (within brain regions that showed significant attentional modulation for HA > HI; with activation mask from the HA > HI contrast; see Methods).

**Table S3.** Peak anatomical locations, Z values, and MNI coordinates from the whole-brain significant clusters showing areas that were functionally coupled with bilateral FG for ignoring face stimuli relative to encoding face stimuli (FI > FA).

|  | Cluster number | Z value (local maxima) | x | y | z |
| --- | --- | --- | --- | --- | --- |
| Precuneus | 1 (cluster extent: 1318, p = 0.004) | 3.94 | -10 | -74 | 30 |
| Precuneus | 1 | 3.36 | -36 | -42 | 58 |
| Left SPL | 1 | 3.32 | -32 | -54 | 54 |
| Left cuneal cortex | 1 | 3.09 | -4 | -82 | 28 |
|  | 1 | 3.08 | -18 | -86 | 24 |
|  | 1 | 2.97 | -22 | -56 | 44 |
| Precuneus | 2 | 3.75 | 10 | -50 | 56 |
| Postcentral gyrus, precuneus | 2 | 3.45 | 10 | -38 | 58 |
| Postcentral gyrus | 2 (cluster extent: 993, p = 0.02) | 3.32 | 14 | -38 | 54 |
| Right superior LO, precuneus | 2 | 3.02 | 22 | -66 | 38 |
| Precuneus | 2 | 2.9 | 0 | -54 | 60 |
| Right precentral gyrus, | 2 | 2.85 | 8 | -22 | 62 |

**Table S4.** Peak anatomical locations, Z values, and MNI coordinates from the whole-brain significant clusters showing areas that exhibited a positive correlation (across subjects) between the degree of coupling with bilateral FG during selective WM for faces (FA > FI, PPI effect) and the subsequent memory different for faces.

| Cluster Index | Cluster Index | Z | x | y | z |
| --- | --- | --- | --- | --- | --- |
| Right orbital frontal cortex | 2 (cluster extent: 4194, p = 1.19e-7) | 4.1 | 26 | 18 | -18 |
| Right ITG, temporal occipital fusiform cortex | 2 | 3.99 | 46 | -56 | -14 |
| Right inferior LO | 2 | 3.76 | 48 | -76 | -2 |
| Right inferior LO, ITG | 2 | 3.57 | 52 | -60 | -8 |
| Right Heschl's gyrus/transverse temporal gyrus | 2 | 3.48 | 56 | -8 | 2 |
| Right ITG, MTG | 2 | 3.46 | 54 | -54 | -10 |
| Right frontal pole | 1 (cluster extent: 821, p = 0.0475) | 3.39 | 42 | 36 | -18 |
| Medial frontal cortex, frontal pole | 1 | 3.32 | -2 | 54 | -10 |
| Right frontal pole | 1 | 3.19 | 38 | 38 | -14 |
| Ventral medial frontal cortex | 1 | 3.1 | -2 | 40 | -24 |
| Right medial frontal cortex | 1 | 3.07 | 10 | 50 | -10 |
| Right medial frontal cortex | 1 | 3.04 | 6 | 46 | -16 |

**Table S5.**
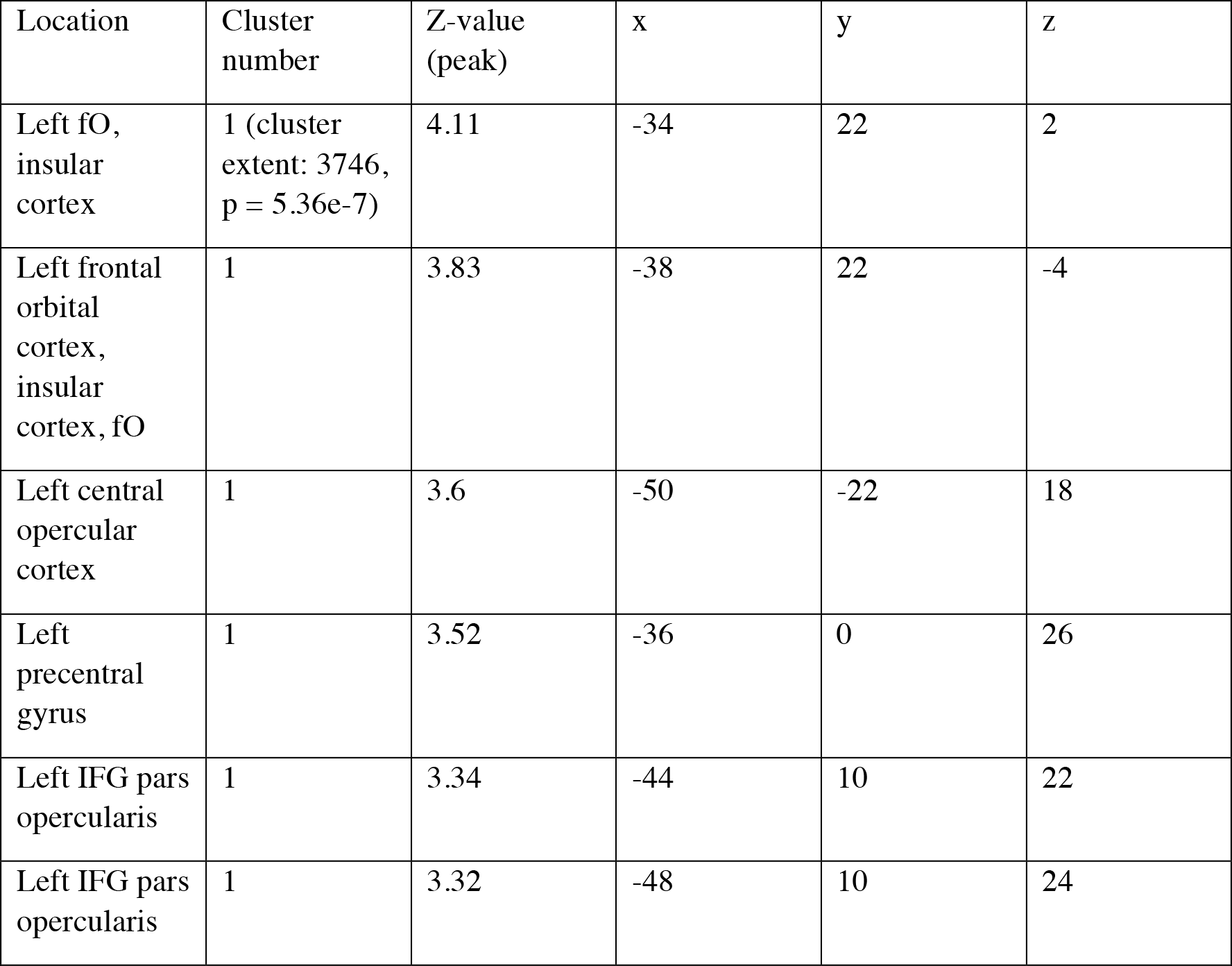
Peak anatomical locations, Z values, and MNI coordinates from the whole-brain significant cluster showing areas that were functionally coupled with bilateral PHG during selective WM for house stimuli (HA > HI).

| Location | Cluster number | Z-value (peak) | x | y | z |
| --- | --- | --- | --- | --- | --- |
| Left fO, insular cortex | 1 (cluster extent: 3746, $p = 5.36e-7$) | 4.11 | -34 | 22 | 2 |
| Left frontal orbital cortex, insular cortex, fO | 1 | 3.83 | -38 | 22 | -4 |
| Left central opercular cortex | 1 | 3.6 | -50 | -22 | 18 |
| Left precentral gyrus | 1 | 3.52 | -36 | 0 | 26 |
| Left IFG pars opercularis | 1 | 3.34 | -44 | 10 | 22 |
| Left IFG pars opercularis | 1 | 3.32 | -48 | 10 | 24 |

